# PMI estimation through metabolomics and potassium analysis on animal vitreous humour

**DOI:** 10.1101/2022.10.20.513043

**Authors:** Emanuela Locci, Matteo Stocchero, Rossella Gottardo, Alberto Chighine, Fabio De-Giorgio, Giulio Ferino, Matteo Nioi, Roberto Demontis, Franco Tagliaro, Ernesto d’Aloja

## Abstract

**Introduction:** The estimation of post-mortem interval remains a major challenge in forensic science. Most of the proposed approaches lack the reliability required to meet the rigorous forensic standards.

**Objectives:** We applied ^1^H NMR metabolomics to estimate PMI on ovine vitreous humour comparing the results with the actual scientific gold standard, namely vitreous potassium concentrations.

**Methods:** Vitreous humour samples were collected in a time frame ranging from 6 to 86 hours after death. Experiments were performed by using ^1^H NMR metabolomics and Ion Capillary Analysis. Data were submitted to multivariate statistical data analysis.

**Results:** A multivariate calibration model was built to estimate PMI based on 47 vitreous humour samples. The model was validated with an independent test set of 24 samples, obtaining a prediction error on the entire range of 6.9 h for PMI<24h, 7.4 h for PMI between 24 and 48h, and 10.3 h for PMI>48 h. Time-related modifications of the ^1^H NMR vitreous metabolomic profile could predict PMI better than potassium up to 48 hours after death, while a combination of the two is better than the single approach for higher PMIs estimation.

**Conclusion:** The present study, although in a proof-of-concept animal model, shows that vitreous metabolomics can be a powerful tool to predict PMI providing a more accurate estimation compared to the widely studied approach based on vitreous potassium concentrations.

## Introduction

Determining the post-mortem interval (PMI) has always been a major challenge for forensic pathologists. Although different methods have been developed over the years [1–10] accurate PMI estimation, highly required in the forensic setting, are still difficult to be obtained.

In consideration of its peculiar anatomy and physiology, the eye has thoroughly been studied for forensic purposes [11–14]. In particular, vitreous humour (VH) has been a biofluid of choice for forensic purposes [15], being investigated through chemical, biochemical, toxicological, and metabolomic approaches to address not only the cause but also the time since death. The post-mortem modifications in VH potassium concentration [K^+^] have probably been the most studied biological parameter to infer the time since death [15].

Recently, multiparametric approaches, such as -omics sciences, appear more suitable to study the multitude of biological phenomena occurring in the post-mortem compared to the traditional methods based on single or few parameters [16–20]. Among them, metabolomics, gradually taking over in forensic research [21, 22], appears suitable to intercept time-related modifications in the early-mid PMI window (up to 72/96 hours). In fact, a growing body of evidence suggests that elapsing time represents the main factor driving metabolome modifications in the post-mortem [23]. Some of the authors have already demonstrated the feasibility of VH metabolome analysis with ^1^H NMR approach [24].

Recently, we applied ^1^H NMR metabolomics to study post-mortem modifications of aqueous humour in an animal model (*Ovis aries*) for PMI estimation [25] using potassium analysis for comparison with an established approach [26]. Through a rigorous statistical evaluation, it was possible to estimate PMI with a prediction error of 100 minutes on the entire time range investigated (24 hours). The results suggested a shared biological phenomenon driving the modifications of both metabolome and [K^+^], being the former able to explain most of the information carried by potassium but showing a greater predictive power in PMI estimation.

The paramount need to investigate higher PMIs fostered the translation of this approach to the VH. To the best of the authors’ knowledge this is the first work in which VH metabolome and potassium are investigated and compared for an accurate estimation of PMI in a controlled animal model.

## Materials and methods

### Samples collection and preparation

Thirty-six heads from young adult female sheep (*Ovis aries*) belonging to the same herd were obtained from a local slaughterhouse after animal sacrifice (for meat consumption) by incision of the jugular vein after electrical stunning. Sheep heads are discarded as waste material, and, consequently, formal approval from the local Ethic Committee was not required for the experimental procedure. All animals were aged between 24 and 48 months and had passed standard controls for food consumption. Sheep heads were rapidly transported to the morgue of the Forensic Science Unit of the University of Cagliari and kept under controlled conditions of humidity (50±5%), and temperature (25±2°C) for all the period of VH collection. Sampling was started at 6 hours after death to allow adequate VH collection for metabolomics, since it was shown that in this time window VH metabolomic composition may be affected by topographical differences of VH sampling [27]. Approximately 1 mL of VH was collected from intact heads using a 5 mL G22 syringe through a single scleral puncture in the lateral canthus. Sampling was carried out at different post-mortem intervals, ranging from 6 to 84 hours. Any eye was sampled once only, to avoid possible contamination. The VH samples taken from the two eyes of the same animal were collected at different times after death. After collection, the VH samples were centrifuged at 13000 g for 5 min to remove any solid debris, mixed with 10 μl of an aqueous solution of sodium azide (NaN_3_, 10% w/w) in order to prevent bacterial growth, and immediately frozen at −80°C. A stratified random selection procedure based on the different ranges of PMI was applied to select 47 VH samples for the training set. The remaining 24 samples were used for the test set. One sample was excluded due to blood contamination.

### VH sample preparation for NMR analysis

Before NMR analysis, samples were thawed and ultrafiltered using a 30 kDa filter unit (Amicon-30kDa; Merck Millipore, Darmstadt, Germany) for 10 min at 13000g and 4°C in order to remove macromolecules and active enzymes. Filters were previously washed out from glycerol by adding 500 μl of distilled water and by centrifuging for 10 min at 10000 rpm at room temperature for 15 times. For the NMR analysis, 250 μl of filtered VH were diluted with 350 μl of a 0.33 M phosphate buffer solution (pH=7.4) in D2O (99,9%, Cambridge Isotope Laboratories Inc, Andover, USA) containing the internal standard sodium 3 (trimethylsilyl)propionate-2,2,3,3,-*d4* (TSP, 98 atom % D, Sigma-Aldrich, Milan) at a 0.75 mM final concentration, and transferred into a 5 mm NMR tube. A volume of 650 μl of the final solution was then transferred into a 5 mm NMR tube.

### ^1^H NMR experiments and data processing

^1^H NMR experiments were carried out on a Varian UNITY INOVA 500 spectrometer (Agilent Technologies, CA, USA) operating at 499.839 MHz. Spectra were acquired at 300K using the standard 1D NOESY pulse sequence for water suppression with a mixing time of 1 ms and a recycle time of 21.5 s. Spectra were recorded with a spectral width of 6000 Hz, a 90° pulse, and 128 scans. Spectra were processed using MestReNova software (Version 9.0, Mestrelab Research S.L.). Prior to Fourier transformation the free induction decays (FID) were multiplied by an exponential weighting function equivalent to a line broadening of 0.5 Hz and zero-filled to 128K. All spectra were phased, baseline corrected, and referenced to TSP at 0.00 ppm. The spectral region 0.80 – 9.00 ppm was segmented into buckets of 0.02 ppm width. The integrated area within each bin was normalized to a constant sum of 100 for each spectrum. The region containing the residual water resonance was excluded before integration. The assignment of the metabolites in the ^1^H NMR spectra was performed based on literature data [24, 27], HMDB database (http://www.hmdb.ca) [28], Chenomx NMR Suite 8.2 Library (Chenomx Inc., Edmonton, Canada), and comparison with spectra of standard compounds recorded using the same experimental conditions. Moreover, using the Chenomx NMR Suite Profiler tool a set of 52 quantified metabolites was obtained (Supplementary Table 1). The final data set was exported as a text file for multivariate statistical data analysis. Mean centering was applied prior to performing data analysis.

### CIA experiments for potassium determination

#### Standards and Chemicals

Standard solutions of potassium (K^+^) and barium (Ba^2+^) were prepared from AnalaR salts (KCl and BaCl_2_) (Merck, Darmstadt, Germany). 18-crown-6 ether (99% pure) and α-hydroxybutyric acid (HIBA) (99% pure) were obtained from Aldrich (Milan, Italy); imidazole (99% pure) and glacial acetic acid was obtained from Sigma (St. Louis, MO, USA). All chemicals were of analytical-reagent grade. Ultrapure water was obtained by an ELGA VEOLIA (Lane End, High Wycombe, UK) water purification system.

#### Instrumentation

All experiments were performed using a P/ACE MDQ Capillary Electrophoresis System (Beckman, Fullerton, CA, USA) equipped with a UV filter detector set at 214 nm wavelength, with indirect detection. In all the experiments, untreated fused-silica capillaries (75 μm I.D., 50 cm effective length; Beckman) were used. The capillary was thermostated at 25ºC. Beckman P/ACE Station (version 8.0) software was used for instrument control, data acquisition and processing. Separations were performed as previously described [29]. Briefly, the running buffer was composed of 5 mM imidazole, 6 mM HIBA and 5 mM 18-crown-6 ether adjusted to pH 4.5 with acetic acid. Constant voltage runs were performed in all experiments by applying a field of 500 V/cm. The analytes were injected at the anodic end of the capillary at 0.5 psi for 10 s. Prior to analysis, all samples were diluted 1:20 with a 40 μg/mL solution of BaCl_2_ (internal standard). Between consecutive runs the capillary was washed with water (3 min) and then with the running buffer (2 min).

### Statistical data analysis

Exploratory data analysis was performed by Principal Component Analysis (PCA) to discover outliers and specific trends in the data. Thus, supervised data analysis based on Projection to Latent Structure regression (PLS) [30] was applied to evaluate the effects of PMI on the metabolomic profiles of the collected samples.

Since PMI showed a non-linear behaviour with respect to the metabolite content of VH, two different regression approaches were applied.

The first approach considered PMI as an ordinal variable after dividing the PMI-window into intervals. Because the metabolite concentrations measured in VH were strongly correlated and the number of training samples was smaller than the number of predictors, standard regression methods developed for ordinal data such as ordered logistic regression could not be applied. For this reason, we developed a new method based on PLS. The method is a three-step procedure where in the first step the levels of the ordinal variable are used as classes to drive a PLS for classification (PLS2-C) [31] model whose score components are used as predictors in the second step where post-transformation of PLS (pfPLS2) [30] is used to model the ordinal variable represented by means of the rank of its levels. As a result, an order between the levels of the ordinal variable is introduced. After these two steps, the latent variable explaining the ordinal variable is calculated. To transform the continuous latent variable into a categorical variable, a Naïve Bayes classifier is built. Since PLS-techniques are robust in the presence of predictor matrices that are not full-column rank matrices, the method is suitable for metabolomics investigations. Repeated N-fold cross-validation was applied in model optimization to estimate the number of PLS components to use for PLS2-C and ptPLS2, and randomization test to assess the reliability of the obtained model. A suitable cost function based on the misclassification error was minimised during cross-validation. Selectivity ratio (SR) [32] based on the predictive component of the ptPLS2 model was calculated to discover the metabolites mainly associated to PMI.

The second approach was developed to directly estimate PMI by regression. Since PMI has only positive values, we considered the logit-transformation to map PMI into a new space where regression was performed. Specifically, PMI was linearly mapped into the closed interval [ε_1_,ε_2_], being ε_1_ and ε_2_ two model parameters in ]0,1[ with ε_2_>ε_1_ to be estimated from the data, and the images ε of PMI in [ε_1_,ε_2_] were logit-transformed to obtain the response to be submitted to PLS2 regression. PMI was, then, predicted applying the PLS2 model on the metabolic profile and using the logistic function to obtain the image ε of PMI in ]0,1[. That image was linearly transformed into PMI in the range [0,L] where L is the maximum value assumed for the prediction of PMI. The procedure assured that the predicted values were positive. Moreover, non-linearity could be introduced modifying the model parameters ε_1_ and ε_2_. Indeed, assuming ε_1_ and ε_2_ close to 0.5 the model was linear whereas non-linearity was introduced for ε_1_ and ε_2_ moving towards 0 or 1. Parameters ε_1_ and ε_2_ and the number of PLS2-components were optimized by repeated N-fold cross-validation maximizing the R^2^ calculated by cross-validation, i.e. Q^2^. Randomization test was applied to test the reliability of the model. Model interpretation in terms of single predictors was performed calculating the SR parameter.

PCA was performed by SIMCA 17 (Sartorius Stedim Data Analytics AB) whereas suitable in-house functions implemented by R 4.0.4 platform (R Foundation for Statistical Computing) were built to perform PLS-based data analysis.

## Results

### CIA [K^+^] determination

Potassium concentrations were determined in 71 VH samples collected at different PMI, ranging from 6 to 84 hours. The potassium concentrations in the analyzed samples ranged from 7.60 to 39.20 mM.

### VH metabolomic profile vs. PMI

VH samples (n=71) collected at different PMI’s, ranging from 6 to 84 hours, were analysed by ^1^H NMR. The 47 samples selected for the training set and the remaining 24 samples used as test set covered the entire investigated PMI window. Outlier detection was performed considering the centered binned data and applying the T2 Hotelling test and the Q-residual distance test. Assuming α=0.05, no outliers were detected.

Exploratory data analysis by PCA applied to the autoscaled set of 52 quantified metabolites generated a model with 2 principal components, R^2^=0.488 and Q^2^=0.363. The score scatter plot and the loading plot are reported in Fig. 1. Colouring the samples in the score scatter plot (Fig. 1A) according to PMI shows a PMI-related trajectory of samples from the upper right-hand corner to the bottom left-hand corner, proving that the data variation of the quantified metabolites included information about PMI. Specifically, the investigation of the loading plot (Fig. 1B) allowed the identification of single metabolites mainly associated to PMI, as indicated in the caption of Fig. 1.

**Fig. 1.**
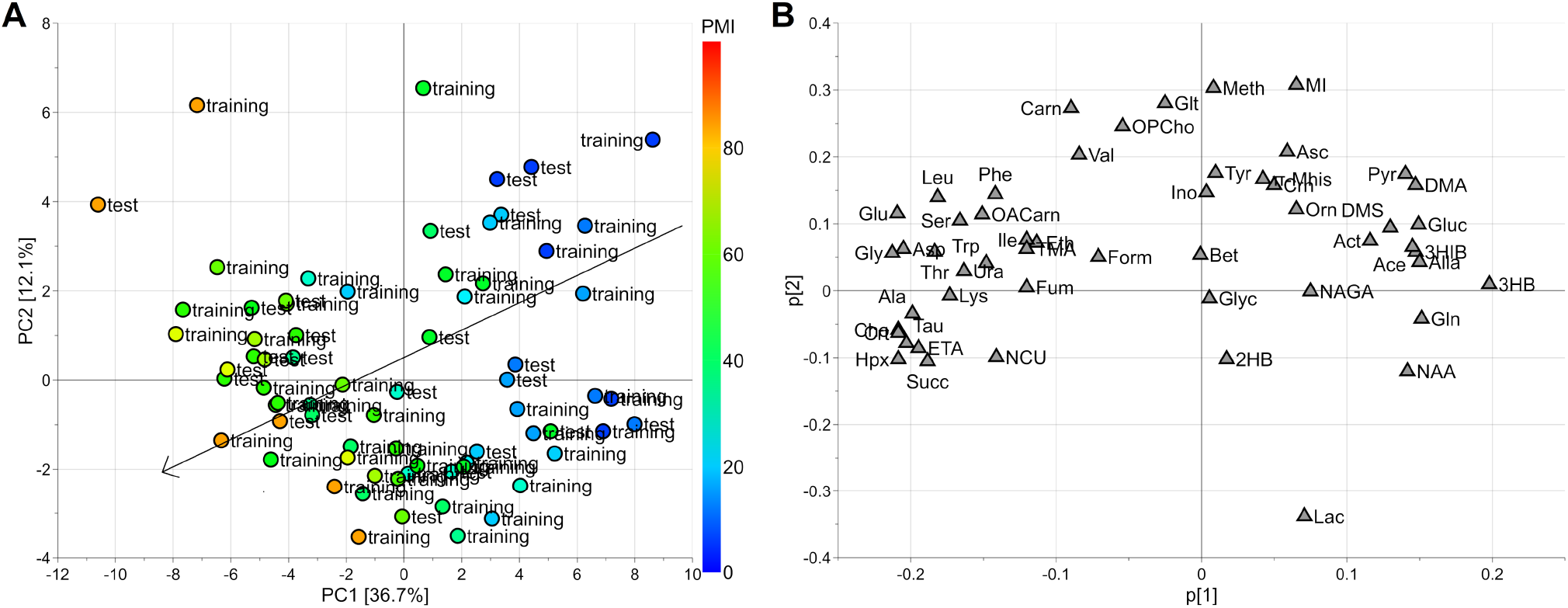
PCA model: PMI increases from the upper right-hand corner to the bottom left-hand corner (panel A). Investigating the loading plot (panel B), the decreasing of glucose, pyruvate, 3-OH-butyrate with the increasing of PMI is observed, while late PMI samples (48-84 hours) are characterized by higher levels of taurine, choline, creatine, hypoxanthine, ethanolamine, and succinate. Samples of the training set are indicated as “training” whereas samples of the test set as “test”.

After exploratory data analysis, regression modelling was applied. Firstly, the PMI temporal window was split into three intervals corresponding to PMI less than 24 hours (interval A), PMI from 24 to 48 hours (interval B) and PMI greater than 48 hours (interval C), respectively, and an ordinal regression model based on PLS was built to predict the interval of PMI given the quantified metabolites. Samples resulted to be balanced with respect to the PMI intervals: interval A included 15 training samples and 7 test samples, interval B included 14 training and 8 test samples whereas 18 training and 9 test samples were included in interval C. Autoscaling the data, the model showed 3 components for the PLS2-C part and 2 components for the ptPLS2 part. The model passed the randomization test (1000 random permutation) assuming α=0.05. The results in calculation, cross-validation (20 repetitions of 5-fold cross-validation) and predicting the test set have been reported as confusion matrices in Fig. 2. It is worth noting that misclassification errors occurred only in contiguous intervals. In particular, when the test set is used, only two samples belonging to interval B were misclassified, one being predicted to belong to interval A and the other to interval C. Assuming α=0.05, we found that alanine, creatine, succinate, glycine, hypoxanthine, choline, ethanolamine, glutamate, taurine, and 3-hydroxybutyrate resulted to be significantly related to PMI on the basis of SR. Only 3-hydroxybutyrate decreased with PMI while the other relevant metabolites were positively correlated to PMI.

**Fig. 2.**
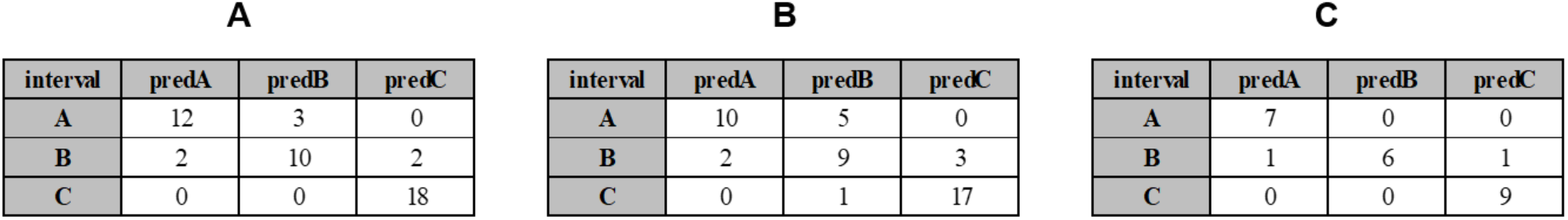
Regression model for ordinal data: confusion matrices obtained calculating the training set (A), performing cross-validation (B) and predicting the test set (C).

Thus, PMI was directly modelled using the new PLS approach developed for positive response and non-linear regression. The dataset of the quantified metabolites was log-transformed and autoscaled. The model showed 3 components, ε_1_=0.02, ε_2_=0.10, R2=0.941 (p<0.001) and Q^2^=0.852 (p<0.001). The model passed the randomization test (1000 random permutation) assuming α=0.05. The root mean square error (RMSE) in calculation was 5.6 hours, RMSE by 20-repeated 5-fold cross-validation was 8.9 hours and RMSE obtained predicting the test set was 8.5 hours. Considering the intervals defined in the case of ordinal regression, the RMSEs estimated using the test set were 6.9 hours for PMI<24 hours, 7.4 hours for PMI between 24 and 48 hours and 10.3 hours for PMI>48 hours. Assuming α=0.05, the analysis of the SR spectrum allowed us to discover threonine, alanine, glutamate, and glycine that are positively correlated to PMI, and glucose and 3-hydroxybutyrate, that are negatively correlated to PMI, as significantly relevant. The results of the interpretation of the regression models are summarised in Fig. 3.

**Fig. 3.**
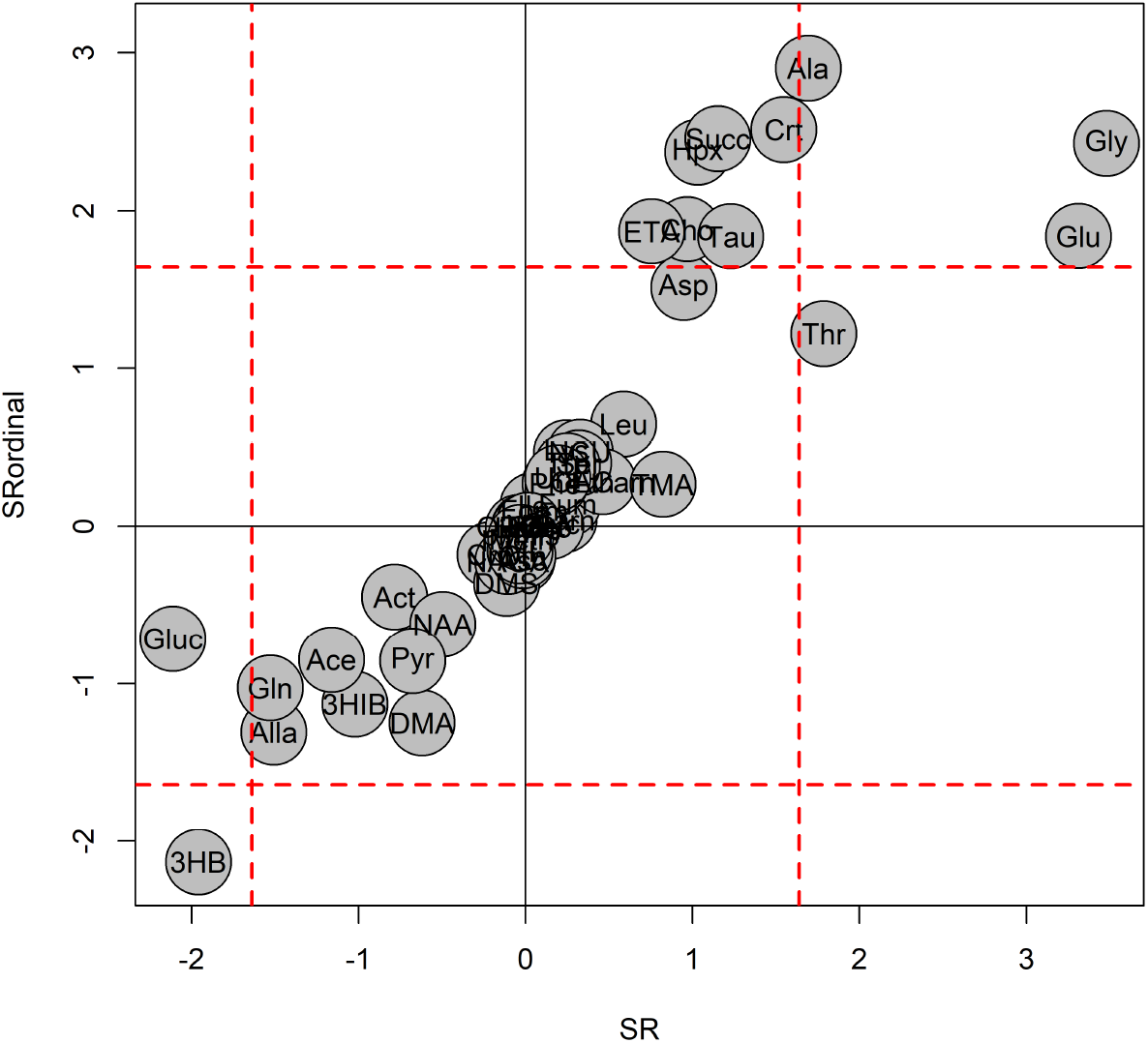
The SR obtained for the ordinal regression model (SRordinal) and that of the regression model (SR) are reported in the same plot; both the ordinal and the regression models discovered 3-hydroxybutyrate, negatively correlated to PMI, and alanine, glutamate, and glycine, positively correlated to PMI, as significantly relevant in predicting PMI (the profiles of these metabolites are reported in Fig. S2 of Supplementary Materials). The SR-values have been multiplied by the sign of the Pearson correlation coefficient calculated between PMI and metabolite concentration; dashed red lines indicate the thresholds of SR at level α=0.05.

### [K^+^] vs. PMI

The relationship between vitreous potassium concentration [K^+^] and PMI was investigated by linear regression. The model showed R^2^=0.582 (p<0.001), Q^2^=0.543 (p<0.001), RMSE in calculation equal to 14.7 hours, RMSE estimated by 20-repeated 5-fold cross-validation equal to 15.4 hours and RMSE in prediction equal to 11.5 hours. Specifically, the RMSEs estimated using the test set were 14.2 hours for PMI<24 hours, 8.8 hours for PMI between 24 and 48 hours and 11.2 hours for PMI>48 hours. In Fig. 4 we have reported in the same plot [K^+^] vs PMI.

**Fig. 4.**
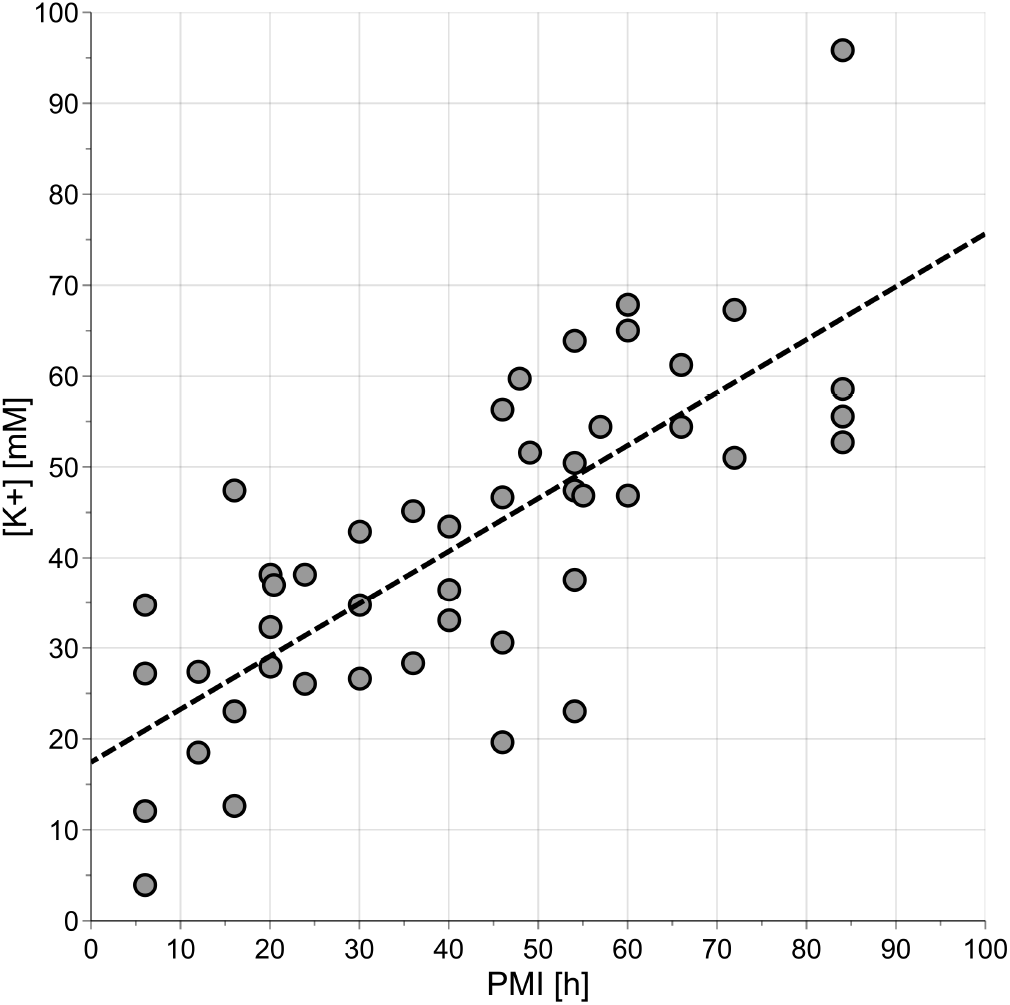
Potassium concentration ([K^+^]) vs. PMI; the dashed line indicates the regression line estimated by linear regression.

The prediction errors resulted to be worse than those obtained considering the quantified metabolites. This is particularly relevant for PMI less than 24 hours. However, potassium concentration may help in predicting PMI if considered together with metabolite concentration. As a proof-of-principle, we investigated the relationship between potassium concentration and quantified metabolites by PLS2 regression. Metabolic data were log-transformed and autoscaled. The PLS2 model showed 1 component, R^2^=0.631 (p<0.001) and Q^2^=0.535 (p<0.001). The metabolites that better explain [K^+^] were discovered by SR (Fig. 5).

**Fig. 5.**
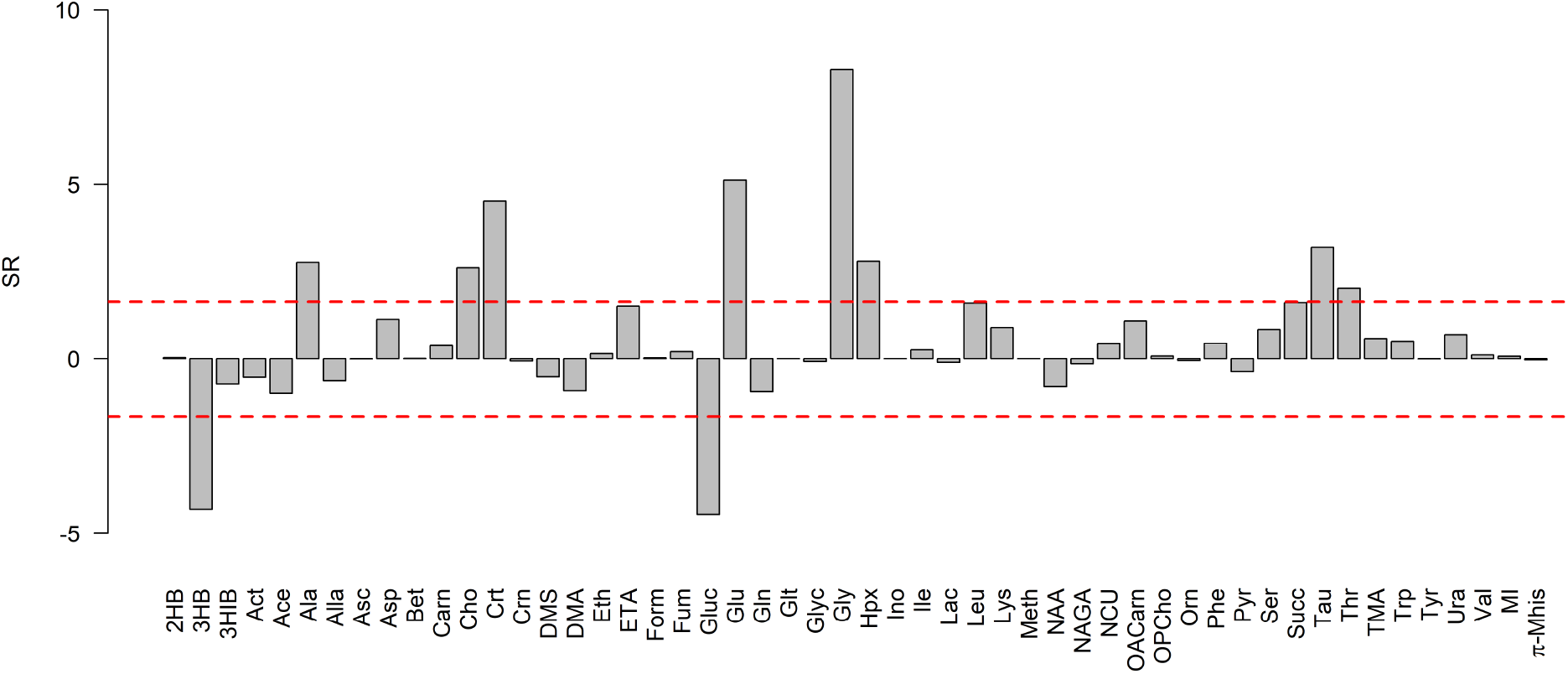
Potassium concentration vs. quantified metabolites: SR plot; threonine, choline, alanine, hypoxanthine, taurine, creatine, glutamate, and glycine, positively correlated to [K^+^], and glucose and 3-hydroxybutyrate, negatively correlated to [K^+^], were discovered as relevant. The SR-values have been multiplied by the sign of the Pearson correlation coefficient calculated between [K^+^] and metabolite concentration; dashed red lines indicate the thresholds of SR at level α=0.05.

Considering the unexplained variance of [K^+^] and modelling PMI by linear regression, we obtained a model with R^2^=0.022 (p=0.315) and Q^2^=−0.097 (p=0.760) that resulted to be unreliable. We could conclude that most of the data variation of the potassium concentration can be explained by the quantified metabolites and that the unexplained variance is not related to PMI. Therefore, the use of potassium concentration together with metabolite concentration may not lead to a relevant improvement in the modelling of PMI with respect to the use of the quantified metabolites alone.

### VH metabolites and [K^+^] vs. PMI

The 52 quantified metabolites and the potassium concentration were considered to model PMI using the non-linear PLS regression approach. Metabolic data were log-transformed, and the potassium concentration was included as [K^+^]^γ^ being the power γ estimated from the data. The parameters γ, ε_1_, ε_2_ and the number of components of the PLS2 model were optimized maximizing Q^2^ calculated by 20-repeated 5-fold cross-validation. The best model showed 2 components, γ=3, ε_1_=0.015, ε_2_=0.10, R^2^=0.942 (p<0.001) and Q^2^=0.871 (p<0.001). The model passed the randomization test (1000 random permutations) assuming α=0.05. The RMSE in calculation was 5.5 hours, the RMSE in cross-validation 8.4 hours, and the RMSE in prediction 7.4 hours. Specifically, the errors estimated using the test set were 5.7 hours for PMI<24 hours, 7.4 hours for PMI between 24 and 48 hours and 8.4 hours for PMI>48 hours. In table 1 we have summarised the results of the obtained models.

**Table 1.** Summary of the results of the regression models: RMSEC is the RMSE in calculation, RMSECV is RMSE calculated by cross-validation and RMSEP is RMSE calculated predicting the test set. Errors are expressed in hours. The PMI predicted for the test set is reported in Fig. S1 of Supplementary Materials.

| model | RMSEC | RMSECV | RMSEP | RMSEP<br>PMI≤24h | RMSEP<br>24h<PMI≤48h | RMSEP<br>PMI>48h |
| --- | --- | --- | --- | --- | --- | --- |
| metabolites | 5.6 | 8.9 | 8.5 | 6.9 | 7.4 | 10.3 |
| [K<sup>+</sup>] | 14.7 | 15.4 | 11.5 | 14.2 | 8.8 | 11.2 |
| [K<sup>+</sup>] & metabolites | 5.5 | 8.4 | 7.4 | 5.7 | 7.4 | 8.4 |

As expected, the improvement obtained including potassium concentration in the set of the predictors is weak and mainly concerns PMI after 48 hours. Moreover, [K^+^] is not significantly relevant if one considers SR and α=0.05.

## Discussion

VH has been long established as a biofluid of choice in forensic post-mortem investigation, with particular attention to PMI estimation. The work here presented is part of a wider project aimed at testing new methods for PMI estimation over different temporal windows in animal models. The rationale of our approach was based on the specific choice of biological matrices appropriate for the study of different PMI ranges. In particular, AH was successfully tested to accurately establish PMI up to 24 hours after death, with a new approach combining ^1^H NMR metabolomic and potassium analysis [26]. Since AH cannot be usually sampled at higher PMIs, the approach was translated to VH which has been traditionally used for this purpose [13, 15].

The observed post-mortem VH metabolome modifications were exploited to build a predictive model for PMI and was compared with the model based on VH potassium obtained in the same dataset, being the latter the actual scientific gold standard for PMI estimation. In the hypothesis of investigating two different biological phenomena, the combined use of the two experimental approaches was also tested.

Results show that VH metabolomic composition is gradually modified both qualitatively and quantitatively with increasing PMIs. The best regression model for PMI estimation obtained using VH quantified metabolites is a non-linear model characterized by a high predictive ability; indeed, when tested on an independent set of VH samples, it shows an error in prediction of 8.5 hours over the entire PMI range (80 hours ranging from the 6^th^ to the 86^th^ hour after death). When three selected PMI ranges are considered (<24, 24-48, >48 hours) the errors in prediction increase with increasing PMI. This is foreseeable as both endogenous and exogenous inferences may cause a progressively increasing biological complexity, yielding to less detectable phenomena.

Another approach using sample classification over three selected PMI ranges appeared very robust, allowing 22 out of 24 samples to be correctly assigned to the proper range. Interestingly, the two misclassified samples belong to the intermediate range, and they occur between contiguous classes. This is consistent with the fact that modifications related to post-mortem interval are continuous, progressive, and developing phenomena.

Focusing on qualitative features of the VH metabolome, glycine, glutamate, alanine, creatine, choline, succinate, hypoxanthine, taurine, threonine, and ethanolamine showed a strong positive correlation with PMI, whereas 3-hydroxybutyrate and glucose were negatively correlated. The observed metabolomic modifications showed a linear behaviour over the entire PMI range, as no specific features can be identified in the three selected ranges.

The best PMI regression model obtained with VH potassium concentrations displays a lower predictive ability compared to metabolomics. The errors in predictions resulted higher over the entire PMI range mainly due to the error in the first 24 hours, which tends to improve in later PMIs, reaching a predictivity with comparable metabolomics. This is consistent with literature data, suggesting human VH potassium as a valuable PMI estimation tool above 24 hours after death (PMI ranging from 2 to 110 hours) [29].

As we previously reported in the AH model, a multivariate metabolomic approach better describes the multifactorial post-mortem phenomenon compared to the best univariate model (i.e., based on a single metabolite taken from the profile) – [23, 26].

In the hypothesis that the two approaches inherently carry different or complementary biological information, we used a combined model. While in early and middle PMI ranges the predictive ability is comparable to the one obtained by the sole metabolome, potassium contribution becomes noticeable in the longest PMI range (> 48 hours). This can be explained by the fact that all the potassium variance is already included in the metabolome modifications up to 48 hours. At higher PMIs metabolome is likely influenced by exogenous bacterial contribution whereas potassium might still reflect the endogenous phenomenon.

This study has several limitations. The investigation was conducted on an animal model, a highly homogeneous and controlled experimental setup being mandatory to precisely estimate PMIs. The translation to human cases represents the following step and should include real-life scenario, different environmental conditions, and multiple causes of death. Furthermore, VH was sampled from isolated sheep heads leading to a possible exogenous bacterial interference on both metabolome and potassium, although the model seems to be reliable and robust. Lastly, VH samples were delivered from the University of Cagliari, where the animal experiment was performed, to the University of Verona for potassium analysis; this may have influenced sample stability, and therefore the potassium CIA results, although the behaviour on the investigated range is consistent with literature data.

To the best of our knowledge, this study and the previous one on AH [26] proposed a tool for PMI estimation providing for the first time PMI prediction model externally validated using an independent set of samples. This model validation strategy, which is a specific issue of potassium according to the literature [33], is mandatory to assess interlaboratory reproducibility, a fundamental step toward the implementation in routine casework.

## Authors’ contributions

All authors read and approved the final manuscript.

## Data availability

Datasets generated and/or analysed during the current study are available from the corresponding author on reasonable request.

## Ethics declarations

This article does not contain any studies with human participants performed by any of the authors. As sheep heads represent waste material, there were neither need of an *ad hoc* animal protocol nor associated cost.

