## Supplementary Materials for "PMI estimation through metabolomics and potassium analysis on animal vitreous humour"

**Fig. S1** Values of PMI predicted for the test set (PMI predicted) vs measured values (PMI): predictions from the model obtained considering the quantified metabolites (panel A), the vitreous potassium concentration (panel B) and the combination of quantified metabolites and potassium concentration (panel C); the diagonal is reported in the plots.

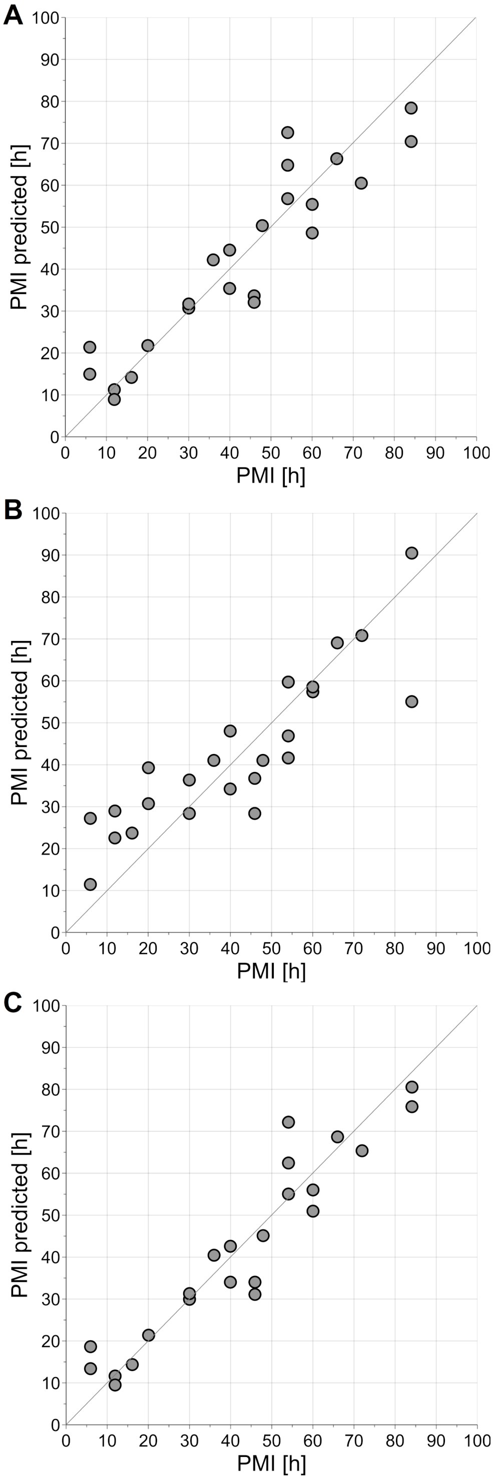

**Fig. S2** Profiles of the significantly relevant metabolites discovered by both ordinal and regression models: 3-hydroxybutyrate (panel A), alanine (panel B), glutamate (panel C) and glycine (panel D); orthogonal scaling has been applied to concentration to have the same scale [-1,1] for all the metabolites; dashed lines indicate the linear regression lines

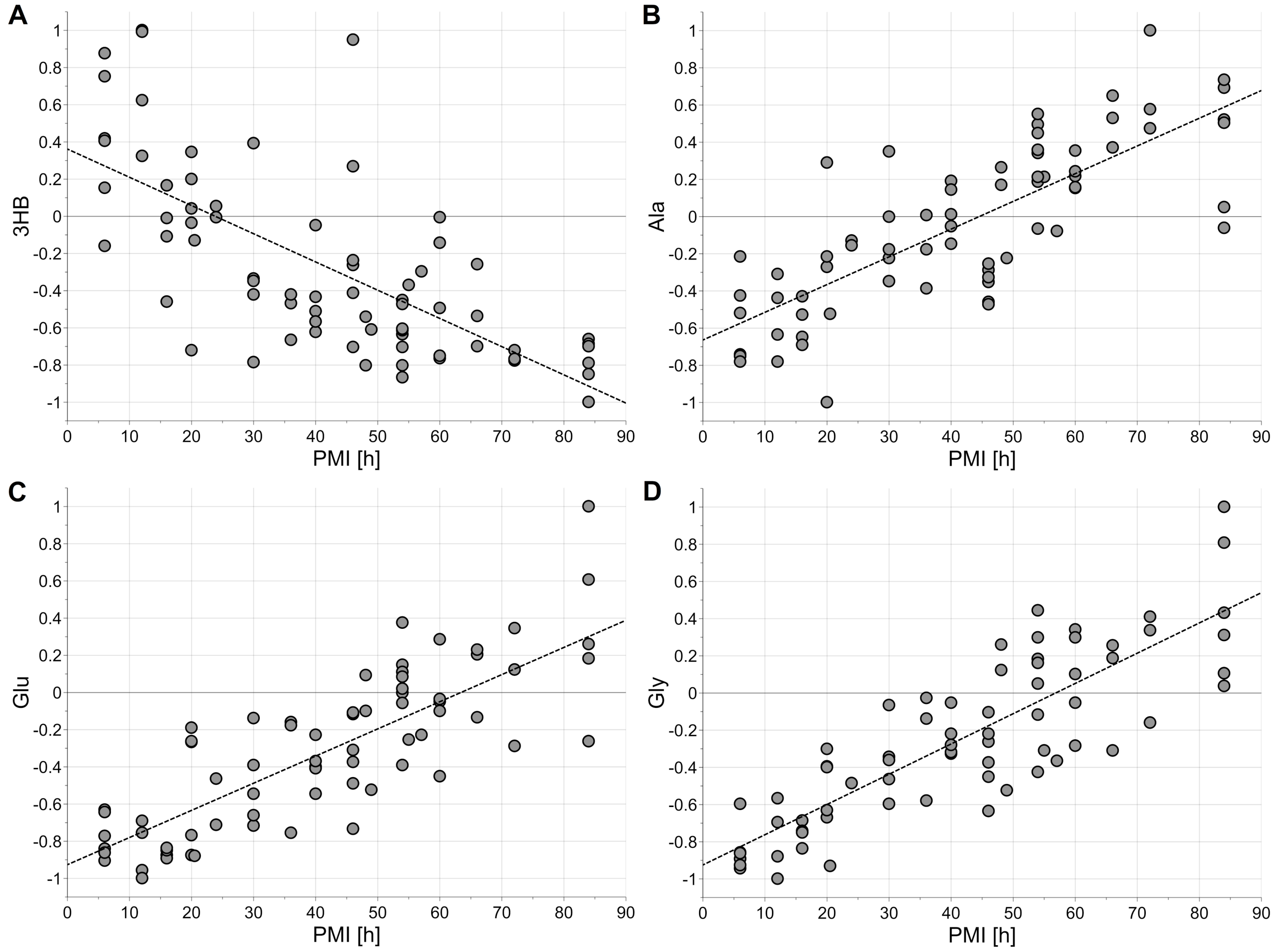
